# Rhizosphere Mysteries: Metabolite Reduction Down-regulated Fungal Diversity and Community Function

**DOI:** 10.1101/2024.03.22.586285

**Authors:** Jichao Li, Zongliang Xu, Tianmei Yang, Jinyu Zhang, Yingmei Zuo

## Abstract

The rhizosphere serves as the primary defense against pathogens, but rhizosphere metabolites can also act as carbon sources and signaling molecules that attract soil borne pathogenic fungi to the destruction of rhizosphere defenses. We propose that whether reducing rhizosphere metabolites improves complex microbial networks defense. Here, we found that reducing rhizosphere metabolites altered fungal community structure more than bacteria, resulting in a downward trend in fungal diversity, soil-borne pathogenic fungal *Fusarium* abundance, and soil microbial community functions, e.g., metabolic functions, enzyme activities, and protein expression. However, the trend is more favorable to plant growth, which might be explained by the combined effect of the upward trend in bacterial diversity in the rhizosphere and bulk soil. Furthermore, we identified biomarkers *Monographella*, *Acremonium*, *Geosmithia*, and *Funneliformis*, which negatively correlated with other differential microbiology, play a competitive role in community member interactions. they optimized the microbial ecology with functions that mobilize soil nutrients, reduce pathogens and soil acidification, and lower phenolic acids. Integrating our findings proposes new avenues for understanding the complex soil rhizosphere mysteries of the critical role of metabolites in “soil environment - microorganisms - metabolites” ecology interactions and provides a design to build synthetic microbial community to enhance defense.

**IMPORTANCE:** While rhizosphere metabolites are known to regulate microorganisms’ composition to enhance plant immunity cooperatively. However, they also have a harmful side, which attracts soil-borne pathogenic fungi to form synergistic damage that inhibits beneficial bacteria, produces autotoxicity, destroys the rhizosphere microbial ecology, and negatively affects soil productivity and plant health. Currently, our planet is experiencing unprecedented anthropogenic-induced changes. Moreover, the complex and dynamic ecological network in the rhizosphere-an important microbial hotspot-is among the most fascinating yet elusive topics in microbial ecology. Whether reduced rhizosphere metabolites improves the microbial ecological networks remains unknown. Our findings revealed that reduced rhizosphere metabolites decrease fungal diversity, microbial community function, and pathogen abundance, while increase bacterial diversity, soil nutrients, pH, and similar factors. It is clear that reduced rhizosphere metabolites is undoubtedly beneficial for plant health and the rhizosphere ecology. Ultimately, This study provided a new comprehensive understanding of how fungi and bacteria assemble and alter in the rhizosphere and bulk soil when reduced rhizosphere metabolites. Understanding the critical role of rhizosphere metabolites in restoring micro-ecological balance will allow us to focus on regulating microbial community metabolism and root exudates, facilitate the discovery of new metabolites and interactions with microorganisms, and harness their the beneficial properties that contribute to rhizosphere microbial community assembly.

---

Root activities, microorganisms, and metabolism influence the rhizosphere, which has previously been reported as the most vital driver In plant growth, resistance, and evolution, referred to as the second plant genome(1). Historically, the microbial communities established in animals, plants, soils, oceans, and virtually every ecological niche on Earth perform vital functions for maintaining the health of the associated ecosystems(2). Microbial communities in the rhizosphere can quickly respond to changes in environmental conditions, particularly during the early stages of crop succession, which is an essential indicator of the study of ecological restoration and community succession(3). Interactions between plants and soil microorganisms that occur in the rhizosphere create microhabitats that result in plant species-specific rhizosphere microbial communities(4). Specifically, plants can change the soil microbiota by secreting bioactive molecules into the rhizosphere(5).

Root exudates typically comprise primary metabolites such as sugars, amino acids, carboxylic acids, and a diverse set of secondary metabolites(6). The root exudate composition reflects the contradictory-concomitantly attractive and repulsive-behaviour of plants towards soil microorganisms. the attractive behaviour is that root exudates used as carbon sources, attracting phytobeneficials that harbor these plant growth-promoting rhizobacteria (PGPR). These PGPRs concurrently perform the functions of nitrogen fixation, nutrient mobilization, growth regulator synthesis, and disease suppression to promote soil microbial production, nutrient availability, and plant productivity(7). They also contain signaling molecules required to establish “plant-microorganisms” interactions. Furthermore, the repulsive behaviour is that plants produce antimicrobial, insecticide, and nematicide compounds to repel pathogens and invaders, as a first line of plant protection, provide biological and physical barriers that can be beneficial to plant health(8). Overall, plants produce root secretions that selectively shape unique soil microbial communities, providing protection and favorable feedback to the plant.

Rhizosphere metabolites also have harmful properties, particularly mediated synergistic damage with soil-borne pathogens disrupting the ecological balance of “plant-soil-microorganism”, causing continuous cropping obstacles(CCOs), where severe in industries that rely on medicinal plants with tuberous roots. Pathogenicity of soil-borne pathogens essentially involves stages of attract (soil to the rhizosphere), colonization (rhizosphere to root surface), and attack (root surface to root)(9). Root secretions can facilitate each of the stages: (i) When root exudates match the preferences of the pathogen, they can support the pathogen’s survival, migration to the rhizosphere, and disease proliferation and serve as a source of carbon and energy. It helps pathogen to reconstruct, stimulates spore germination and mycelial growth, and enhances the expression of chemotaxis and virulence-related genes. Root secretion of ferulic acid enhances the SH3 structure and F-box protein system of the pathogen, regulating the expression of associated genes and increasing the infective capacity of the pathogen(10). Cinnamic acid increased fusaric acid secretion by 247.12% in *Fusarium oxysporum*, which makes plants more vulnerable(11). (ii) restruction of beneficial microbial communities in the rhizosphere.Butyrate acid caused a change in the structure and function of the tobacco rhizosphere microbial community from beneficial to harmful, leading to CCOs(12). Organic acids significantly promoted the production of toxins and H_2_O_2_ secretion by the rhizosphere pathogens of *Radix pseudostellariae* L.and which resulted in attenuated antagonistic activities of PGPR to suppress mycelial growth of the pathogenic fungi(13). (iii) poison root tip tissues, impairing root cell membrane permeability, root vigor, and systemic resistance. The pathogen induces the release of specific root secretions, which increase the virulence and the quantity of chemo-sensory autotoxic substances and induce greater autotoxicity. *Fusarium oxysporum* infection alters the expression of genes related to phenolic acid synthesis in host plants, promoting phenolic synthesis and accumulation in the rhizosphere(14). Phenolic acids significantly reduce the activity of plant defence substances β −1,3-glucanase(15). (iv) change physical and chemical factors such as acidity, fertility, and enzyme activity of the rhizosphere. Plant roots secrete large amounts of organic acids to reduce soil pH, and carrying carboxyl groups of mucilage will also release H^+^ to promote soil acidification, but also to promote pathogenic H+ efflux, further exacerbating the rhizosphere soil acidification. This change creates a suitable acidic environment that inhibits beneficial bacteria and accumulates specific pathogens, disrupting the balance of rhizosphere microbes(16). Integrating, rhizosphere microecological imbalances that common issues focusing on rhizosphere microbial community reconfiguration, autotoxicity, and soil physicochemical properties. However, the key to connecting the three is root secretion.

Rhizosphere microecological imbalances pose a significant challenge to plant immunity and survival. There is currently no answer to completely solving the CCOs and why beneficial microbes do not work (they do in the lab clearly even inhibit the pathogen) due to the complex, dynamic, and puzzling microbial ecological network in the rhizosphere that has yet to be clear, which is among the most fascinating yet elusive topics in microbial ecology.

Despite research on metabolite composition, interactions with microorganisms (in the laboratory), and microbial ecology, there still needs to be a significant knowledge gap about regulating microbial assembly by rhizosphere metabolites, particularly how to achieve beneficial feedback.

Therefore, we hypothesize that reducing root secretions may reduce interactions with pathogens to balance the microbial ecology in CCOs. Could it also result in beneficial microorganisms being unable to survive, affecting the rhizosphere’s ecological function? Hence, We then consider (i) whether reduced rhizosphere metabolites improve the rhizosphere microbial ecology and plant health and (ii) how core microbes shape the rhizosphere resistance when reduced rhizosphere metabolites would provide a design to build synthetic microbial communities next.

Here, we investigate how reducing root secretions (in soil environments) affects the fungi and bacteria alteration in both the rhizosphere and bulk soil and how that affects the soil environment and plant survival. We will identify biomarkers and analyze how they relate to microbial communities, soil nutrients, and autotoxicity. Finally, we propose new avenues for understanding rhizosphere microbial ecology when reducing root secretions and provide a theoretical basis for designing and constructing synthetic microbial communities.

## RESULTS

### Fungal communities highly affected by reduced rhizosphere metabolites than bacteria

To expose how reducing rhizosphere metabolites (in soil environments) affects the fungi and bacteria alteration and optimization strategy of microbial community structure, We employed three strategies that have previously shown efficacy in reducing root secretions, i.e., a biochar strategy, a grass ash strategy, and a corn stover strategy. Subsequently, the design distinguishes between the non- and rhizosphere soil, where less and more influential by root secretion.

However, plant growth and seedling survival rates are different among the strategies, with the biochar strategy having the most favourable phenotype and seedling survival(*P*<0.01)(Fig. S1). Similarly, we found that the microbial community composition displayed a high variability between the fungal community and even between the non- and rhizosphere soil, highlighting the microbiome heterogeneity of soil when reduced rhizosphere metabolites(Fig. 1). Bacterial and fungal communities have discernible differences, the vast majority of bacterial communities have a structural resemblance in response to different strategies(Fig. 1B and D), and fungal communities were most strongly affected by reduced rhizosphere metabolites than bacteria(Fig. 1). In the non-rhizosphere, all the strategies had more fungal richness compared with control group, displaying reduced rhizosphere metabolites limits fungal microbial recruitment to the rhizosphere(Fig. 1C).

**FIG1.**
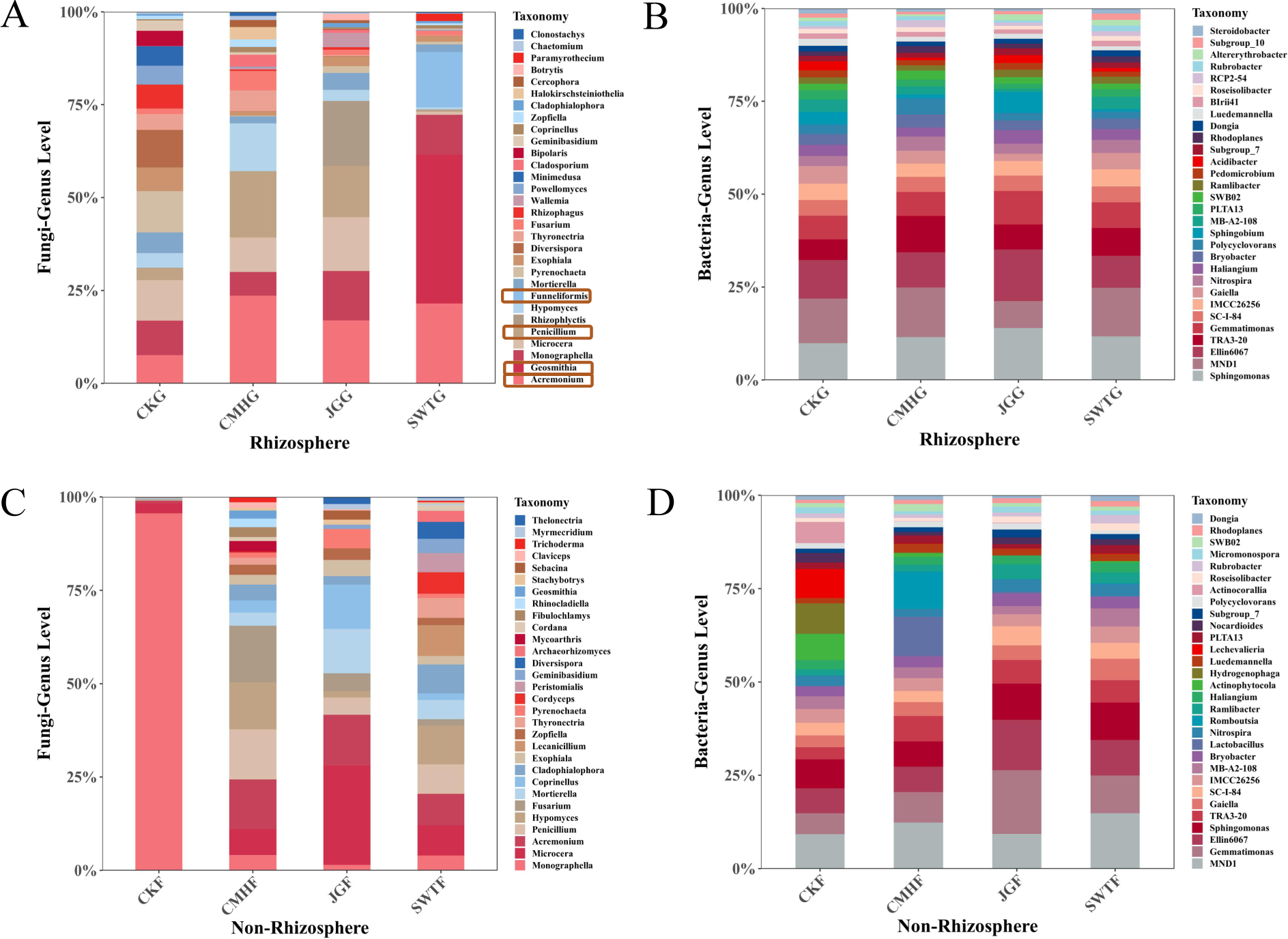
Fungal and bacterial community structure at the genus level. (A) and (B) Fungal and bacterial community structure in the rhizosphere. SWTG (biochar), CMHG (grass ash), JGG (corn stover), and CKG (control), similarly hereinafter. Orange boxes indicate the most changed fungal genera. (C) and (D) Fungal and bacterial community structure in the non-rhizosphere. SWTF (biochar), CMHF (grass ash), JGF (corn stover), and CKF (control), similarly hereinafter. Reducing rhizosphere metabolites had a greater variety of fungal community structures. Conversely, bacterial communities display a similar structural response to various strategies.

The *Acremonium* genus exhibited a highly represented across all three systems, while the richness of *Penicillium* genus was highest in the grass ash strategy(CMHG) and corn stover strategy(JGG). Notably, the biochar strategy(SWTG) revealed a distinct trend of aggregating specific genera to rhizosphere, including *Geosmithia* spp. and *Funneliformis* spp. (Fig. 1A).

### Fungal diversity reversed between the non- and rhizosphere, fungal diversity decreased and bacterial increased in rhizosphere

Following that, we then performed a paired test comparing non- and rhizosphere microbial diversity, investigating variations of rhizosphere microbial in which proximity or distance (more or less influence) of rhizosphere metabolites. The result revealed notable disparities in microbial diversity between the non- and rhizosphere, with a reversal occurring comparing the control, particularly for fungi. The reversal phenomenon was that control showed high fungal diversity in the rhizosphere than non-rhizosphere while reduced rhizosphere metabolites resulted in low in the rhizosphere than non-rhizosphere, i.e., biochar and grass ash strategies (*P*<0.001) (Figs. 2A and B). Bacterial diversity showed no difference between the non- and rhizosphere in the control (Figs. 2C and D), yet there was a statistical difference in the biochar strategy(Fig. 2C).

**FIG2.**
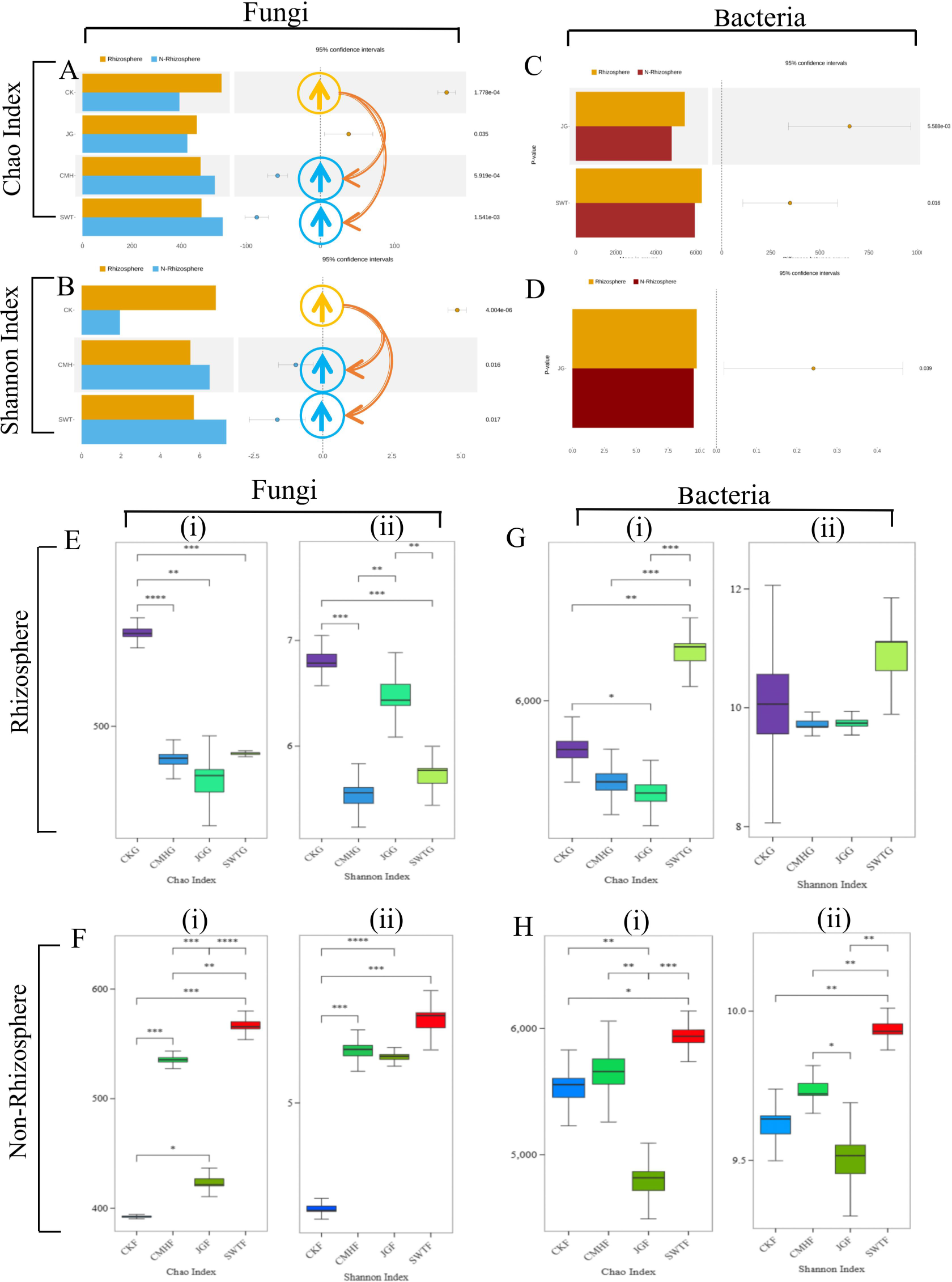
Fungal and bacterial community diversity. (A) and (B) Fungal Chao and Shannon index paired map between non- and rhizosphere (all strategies). It only shows different strategy pairs (same as C and D). The diversity in the rhizosphere overlies the non-rhizosphere with yellow arrows. Non-rhizosphere overlies rhizosphere with blue arrows. Fungal diversity is lower in the rhizosphere with reduced rhizosphere metabolites. (C) and (D) Bacteria Chao and Shannon index paired map between non- and rhizosphere (all strategies). Bacterial diversity is higher in the rhizosphere with reduced rhizosphere metabolites. (E) and (F) Fungal diversity comparison box among all the strategies(i: Chao index, ii: Shannon index. same as F, G, and H). Fungal diversity followed the same trend across strategies. (G) and (H) Bacteria diversity comparison box among all the strategies. Bacterial diversity showed inconsistent trends compared to the control, with a higher diversity with the biochar strategy (SWTG) in both the non- and rhizosphere.

We presented a detailed analysis of the differences in microbial diversity among all the strategies. Fungal diversity followed a similar trend across all strategies on reduce rhizosphere metabolites(Figs. 2E and F), while bacterial diversity varied. We observed a higher diversity with the biochar strategy (SWTG) in both the rhizosphere and non-rhizosphere, a lower with the corn stover strategy (JGG), which was significantly different from the control (*P*<0.05) and biochar strategy (*P*<0.001), and almost no change with the grass ash strategy (CMHG)(Figs. 2Gi and Hi). A focal, the higher bacterial diversity with biochar strategy could be a strategy to improve the microbial environment, which has the most advantageous phenotypic.

Together, Reducing root secretions modified microbial diversity in the rhizosphere and non-rhizosphere, and affecting both proximal rhizosphere and distant regions. Modifying microbial diversity in the rhizosphere, decreasing fungal diversity, and increasing bacterial diversity, which could be a strategy to improve the microbiological environment.

### Reduced rhizosphere metabolites downregulated metabolic pathway, enzyme activity, protein expression

We further investigate functional variations in soil microbial community by tagging gene sequences to predict practical abundance, using the PICRUSt functional prediction, which the prediction accuracy of soil flora is high and can reach more than 85-90%. The result revealed that soil microbial community functions showed a downward trend when reduced rhizosphere metabolites in the rhizosphere, which most down-regulated in KEGGs predicting, protein functions, enzyme activity, and metabolic pathways(Fig. 3, S2).

**Fig3.**
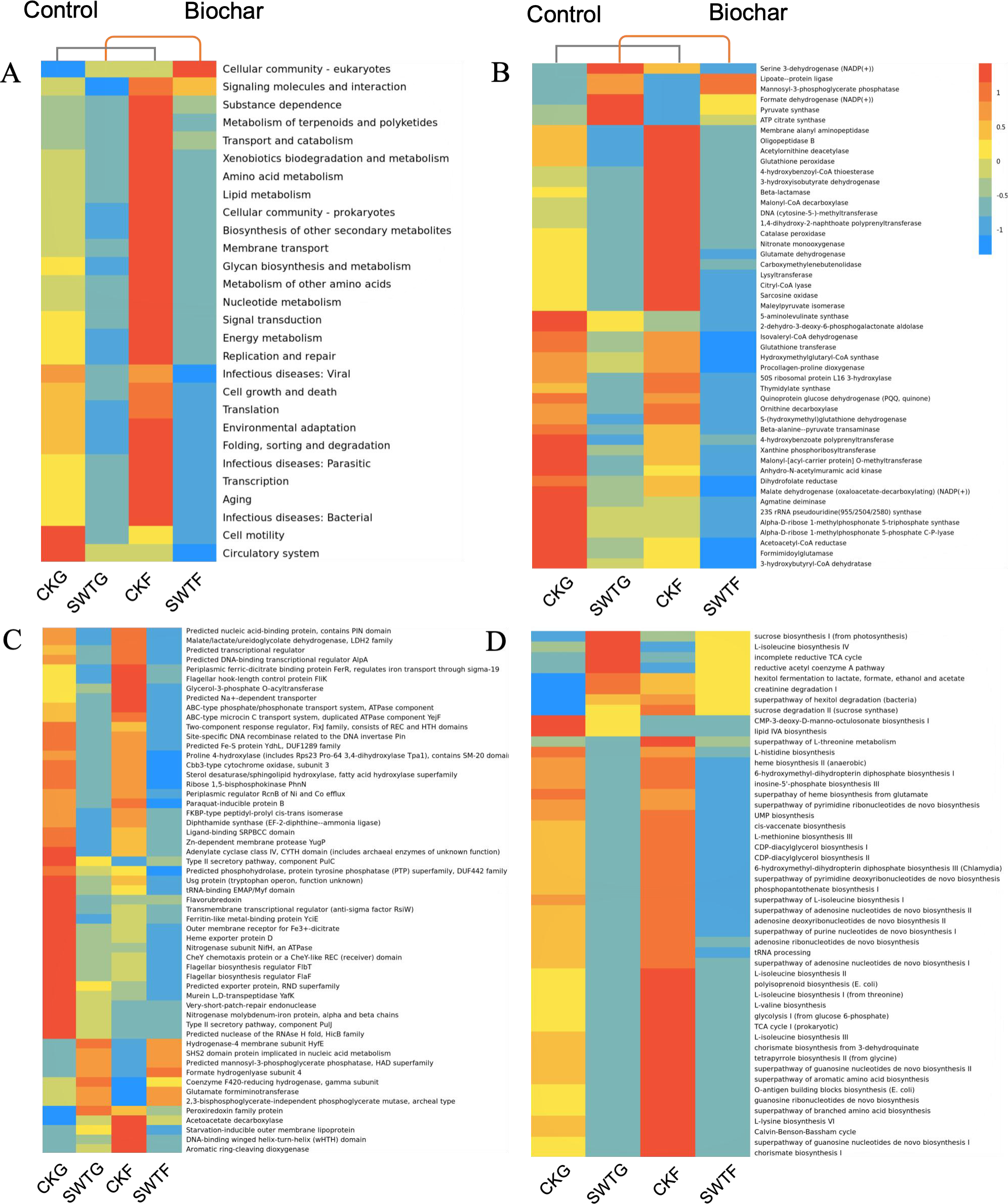
Marker gene sequences predict functional abundance. (A) KEGG prediction difference clustered heatmap(level2). (B) enzyme activity difference clustered heatmap. (C) Protein expression difference clustered heatmap. (D) Metabolic pathway difference clustered heatmap. Soil rhizosphere microbial function were down-regulated when reduced rhizosphere metabolites, including KEGG prediction, protein expression, enzyme activity, and metabolic pathways.

Metabolic functions down-regulated in biochar strategy(SWTG) (Fig. 3, S2). These functions encompassed various processes, such as signaling molecules and interactions, metabolism of terpenoids and polyketides, transport and catabolism, amino acid metabolism, lipid metabolism, biosynthesis of other secondary metabolites, nucleotide metabolism, and energy metabolism(Fig. 3A). Furthermore, cellular processes and genetic information showed signs of down-regulation as well, particularly in processing signaling molecules and interactions, signal transduction, and membrane transport. However, cellular community prokaryotes were down-regulated, while eukaryotes were up-regulated(Fig. 3A).

However, We discovered up-regulation during our investigation, specifically in the form of stimulated enzyme manufacturing and activity, e.g., Formate dehydrogenase (NADP(+)). ATP citrate synthase (NADP(+))(Fig. 3B). Protein expression showed up-regulation, e.g., Hydrogenase-4 membrane subunit HyfE, SHS2 domain protein implicated in nucleic acid metabolism, Formate hydrogenlyase subunit 4, Coenzyme F420-reducing hydrogenase-gamma subunit, Glutamate formiminotransferase, and Perexiredoxin family protein(Fig. 3C). Metabolic pathways that were up-regulated, e.g., sucrose biosynthesis I (from photosynthesis), L-isoleucine brothers IV, the incomplete reductive TCA cycle, and the reductive acetyl coenzyme A pathway(Fig.3D).

Overall, reduced rhizosphere metabolites down-regulated most microbial community functions in the rhizosphere.

### Reduced rhizosphere metabolites accompanied a drop in root autotoxins

We initially considered that biochar adsorbs root secretion as previously reported. To validate our hypothesis, we performed targeted metabolomics on the biochar strategy’s interlayer (biochar-only), determine its ability to absorb metabolomics and which metabolites were most affected.

We present the number of metabolites targeted and quantified to be rhizosphere metabolites, e.g., acids, terpenes, glycosides, and other relevant substances. We found most metabolites were identified and targeted within the biochar strategy’s interlayer. Proves our hypothesis, the biochar strategy had fewer metabolites in the rhizosphere, while had more of them in the biochar strategy’s interlayer(Fig. 4A). It is noteworthy that the metabolites adsorbed to were more phenolic acids with autotoxic effects, e.g., 4-Hydroxybenzoic acid, 4-Hydroxycinnamic acid, Ferulate, Stearic acid, and Salicylic acid, highlight biochar can also reduced root autotoxins(Fig. 4B).

**Fig4.**
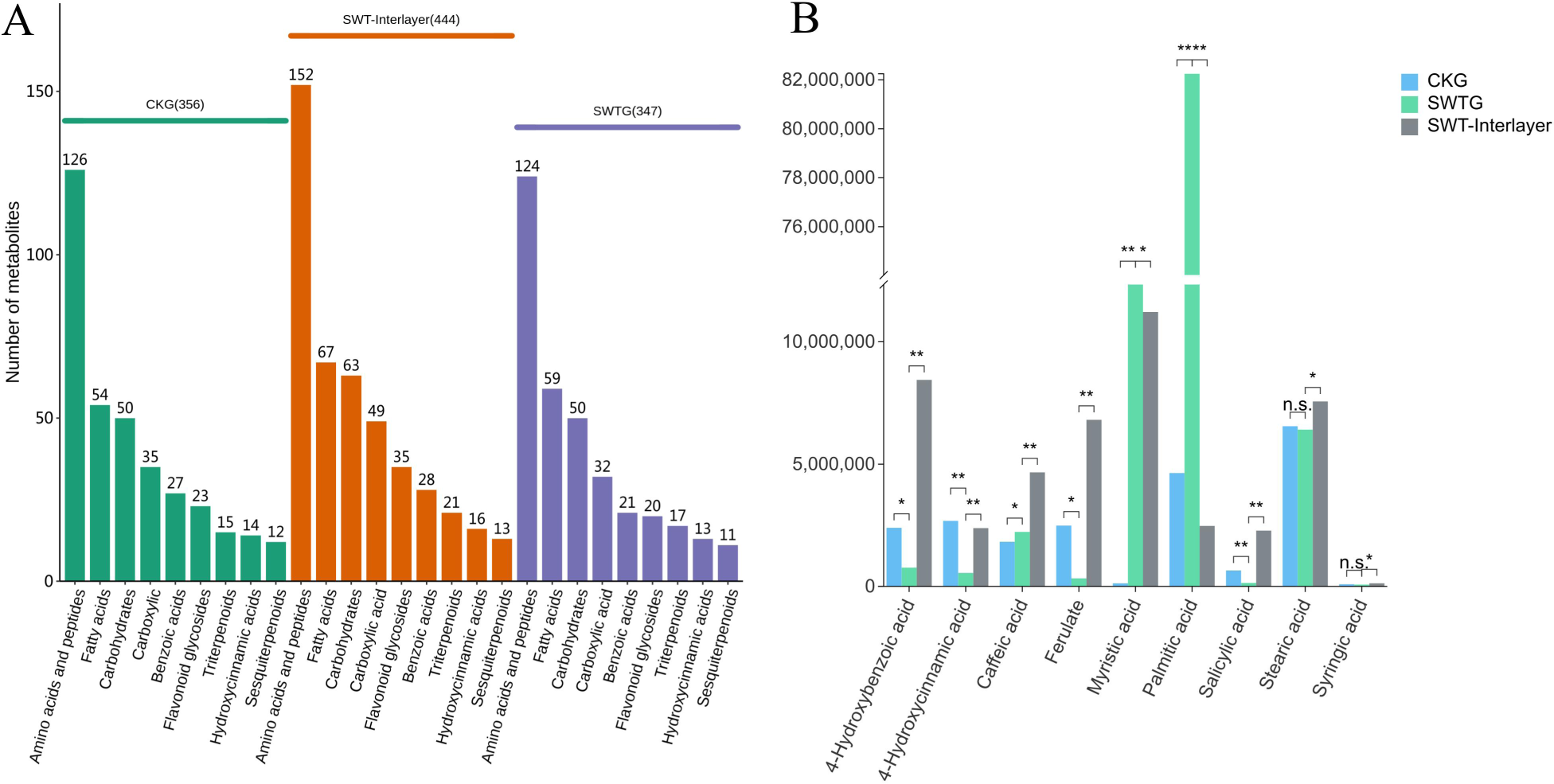
Targeted metabolite count and concentration comparison. (A) Rhizosphere metabolites and precursor counts. The Biochar strategy’s interlayer (biochar-only) contained higher counts of targeted metabolites and precursors. (B) Root autotoxic concentration comparison. Root autotoxins were much lower in the biochar strategy soil rhizosphere while higher in the biochar strategy’s interlayer. Biochar can adsorb root autotoxins and validating that the experiment reduces rhizosphere metabolites in the rhizosphere soil.

Consequently, as previously has been reported, supporting our original prediction that the experiment definitely reduces rhizosphere metabolites and accompanied a drop in root autotoxins in the rhizosphere soil. As a result, This reduction in autotoxins optimizes the rhizosphere microbial ecology by creating a healthier environment for microorganisms to thrive in.

### Fungal biomarkers show competitive roles in microbial interactions

We mainly focus on which functional biomarkers were associated with changes in reduced rhizosphere metabolites to provide design ideas for synthetic microbial community construction, utilizing linear discriminant analysis of effect size (LEfSe), identify biomarkers that exhibited statistically differences between phenotypically superior biochar strategy and controls.

We identified that the fungal biomarkers of phylum Ascomycota and Glomeromycota were most highly associated with functional microbial community. *Monographella*, *Acremonium*, *Geosmithia*, and *Funneliformis* representing the taxa good representation of functional biomarkers.in the biochar strategy(SWTG)(Figs. 5Ac–d).

Further, we found positive correlations between fungal biomarkers in biochar strategy rhizosphere, i.e., between *Monographella*, *Acremonium*, *Geosmithia*, and *Funneliformis*. However, they had negatively correlated with 60% of the top 20 differential fungal microorganisms, uncommonly significantly negatively correlated with the fungal biomarkers in the control rhizosphere.

In addition, The biochar strategy (SWTG) had more bacterial biomarkers than the control (Figs. 5Af, h), which could attract functional bacteria from different species to colonize (Figs. 5Ae–h). The collected bacterial biomarkers besides mainly from *Proteobacteria* and *Actinobacteriota* phylum (Fig. 5Af), also included *Gemmatimonadota* and *Nitrospirota* phylum, which have a wider range of biomarker sources (Fig. 5Af).

**FIG5.**
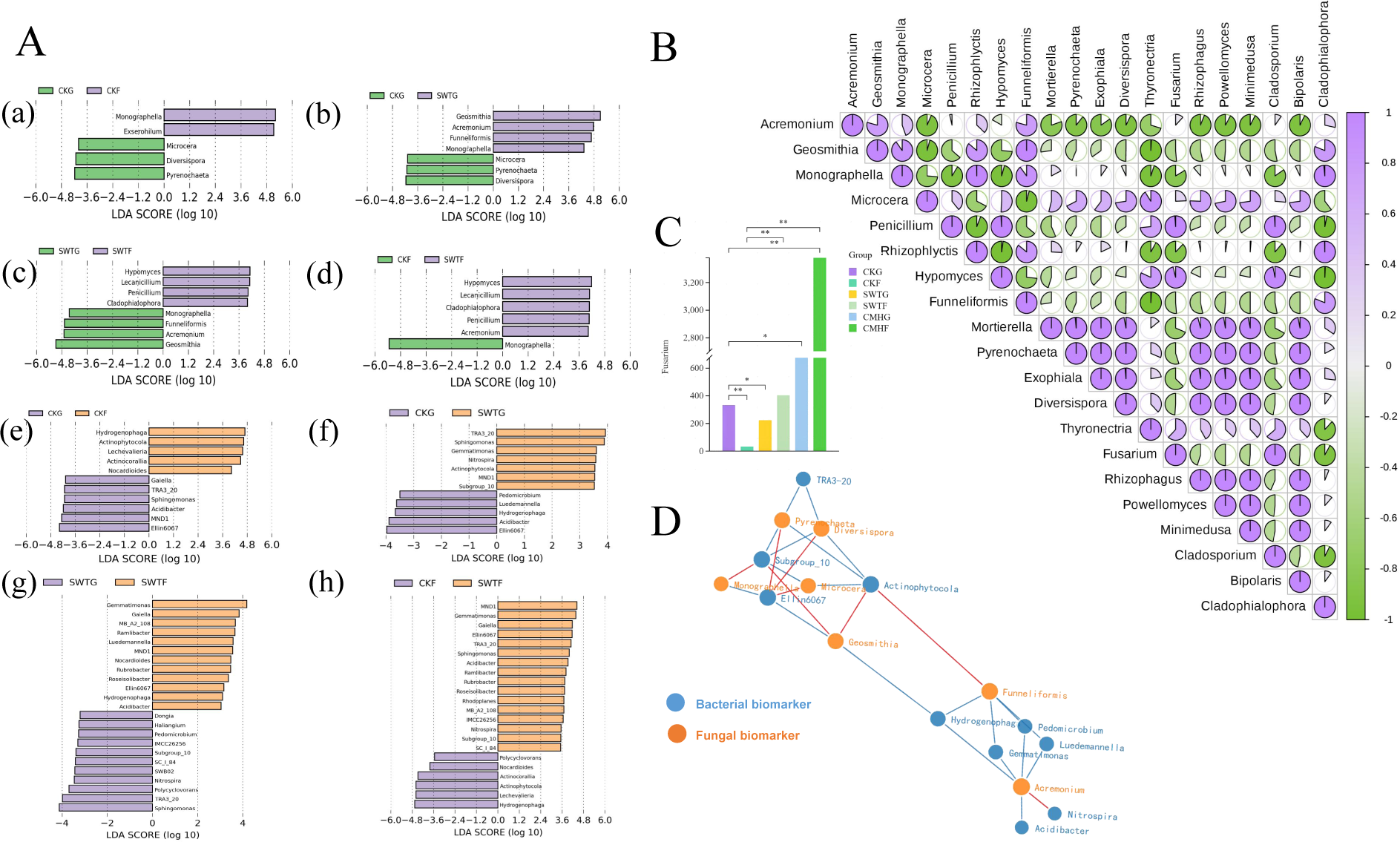
LEfSe and correlation analysis of biomarkers. (A) (a-d) and (e-h) fungal and bacterial microorganisms in LEfSe analysis. Fungal biomarkers in biochar strategy rhizosphere: *Monographella*, *Acremonium*, *Geosmithia*, and *Funneliformis*. (B) Top 20 abundant differential fungus correlation analysis. *Acremonium*, *Geosmithia*, and *Funneliformis* showed a negative correlation with over 60% of the top 20 abundant fungal microbes. (C) Pathogen abundance *Fusarium* spp. histogram. the abundance of *Fusarium* spp. reduced substantially in biochar strategy rhizosphere(SWTG) while rised in grass ash (CMHG). (D) fungal and bacteria biomarkers interomics correlation network. Fungal biomarkers microbiome has a positive correlation with each other but have a negative correlation with bacterial biomarkers.

To gain a better understanding the interactions of biomarkers among community members, construct an interomics correlation network between biomarkers found in fungi and bacteria. Similarly, we found strong negative correlations between the fungal biomarkers, i.e., *Acremonium* and *Funneliformis* with bacterial biomarkers, particularly those found in the control rhizosphere, even stronger than the positive correlation with fungal and bacterial biomarkers within same biochar rhizosphere.

Integrating our findings imply that critical fungal biomarkers with reduced rhizosphere metabolites potentially play a competitive role in microbial interactions within the soil rhizosphere.

We conducted a study on the pathogenic fungus *Fusarium* spp., although not identified as a biomarker, controlling its growth is crucial in restoring the balance of rhizosphere microbial ecology. we focused on changes in the pathogenic fungus *Fusarium* spp. its mitigation is critical to restoring the microbial ecology. Our results showed that the biochar strategy resulted in a statistically decrease in the abundance of *Fusarium* spp. (*P*<0.05) (Fig. 5C), conversely, the grass ash strategy resulted in a significant increase(Fig. 5C), which may explain biochar strategy phenotypic superiority to grass ash.

Soil-borne pathogenic fungal *Fusarium* spp. is a destructive fungus that can disrupt the microbial ecology, and reduced rhizosphere metabolites decreased its abundance, can help maintain a healthy rhizosphere microbial ecology.

### Ecological roles of biomarker microorganisms in the rhizosphere

To better understand the biomarker microorganisms’s role in optimizing the rhizosphere microbial ecology. We also propose a framework on how biomarker microorganisms roles of three major segments, i.e., microbial networks, metabolites, and soil nutrients.

We found fungal biomarkers were positively correlated with soil nutrients and elevated pH but negatively correlated with the pathogenic fungus *Fusarium* spp. and phenolic acid which can be autotoxic to the root system. Conversely, these phenolic acids were negatively associated with soil nutrients and biomarker microorganisms. Therefore, the ecological functions of the fungal biomarkers were more significant for increasing nutrients and decreasing autotoxicity and soil acidification.

The bacteria from the *Acreminium* genus worked synergistically with fungi to perform the same function. Furthermore, NN and pH were the most critical factors in determining the rhizosphere soil environment.

Overall, the biomarker microorganisms played a more significant ecological role in optimizing the rhizosphere environment by efficiently utilizing soil nutrients, defending against pathogens, and participating in the metabolic function of reducing autotoxic metabolites.

**FIG6.**
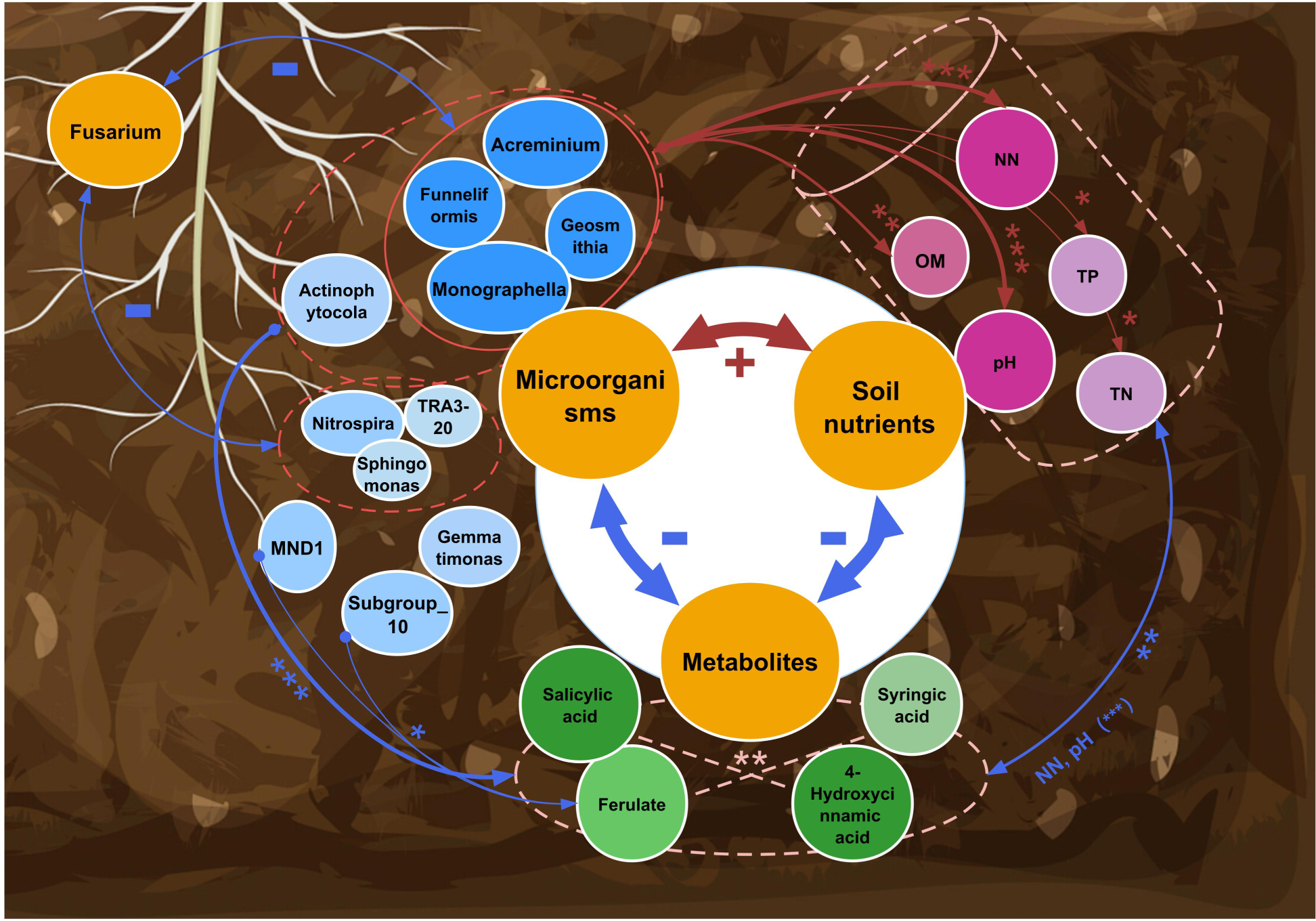
Network interaction maps between microorganisms, metabolites, and soil nutrients. Biomarker microorganisms were positively correlated with soil nutrients and elevated pH but negatively correlated with phenolic acid (*P*<0.001). NN and elevated pH were positively correlated with biomarker microorganisms but negatively correlated with phenolic acid (*P*<0.001). Both fungal and bacterial biomarker microorganisms were negatively correlated with pathogenic *Fusarium*. However, biomarker microorganisms played a more significant role in increasing soil nutrients, reducing phenolic acid and soil acidification, and defending against pathogens.

## DISCUSSION

### Reducing rhizosphere metabolites shifted fungal diversity from more to less in the rhizosphere

This finding suggests that lowered selection and recruitment of microorganisms via root secretions resulted in declined colonization of substrate-preferring pathogens in the rhizosphere. Microbial colonization of plant roots occurs in three stages: attraction by root secretions, colonization, and attack(9). The bulk soil is the seed bank of potential commensals(17). Plant roots release chemical signals that attract beneficial or pathogenic microorganisms, which colonize based on their preferred microbial substrate(18). Genes that encode proteins involved in chemotaxis, flagella assembly, motility, biofilm formation, secretion, and two-component regulatory systems are highly abundant in microorganisms and microbial communities found in the root environment(19), as compared with the bulk soil(20). Therefore, microbial diversity from the non-rhizosphere to the rhizosphere is a process of increasing abundance due to the attraction of root secretions. However, we demonstrated that reducing rhizosphere metabolite caused a shift that became less abundant in the rhizosphere, and higher in the non-rhizosphere, implying that the attraction and maintenance of substrate utilization between microorganisms and roots have been broken.

This shift is helpful as it decreases the process of fungal colonization in the rhizosphere including the pathogen invasion. CCOs soils become dominated by “fungal types” due to the increase in the abundance of soil-borne pathogenic fungi. Pathogens proliferate up the threshold, and plant disease outbreaks begin(21). However, our study found that reducing rhizosphere metabolites results in lower fungal diversity, varied fungal structure, and decreased pathogen *Fusarium*, leading to a weakening in attackers and ecological niches, which would balance microbial ecology. Furthermore, the abundant phenolic acids release by the medicinal plant is the most critical reason why they suffer the severity of CCOs compared to others. phenolic acids are substrate-preferred types of soil-borne pathogenic fungi, attracting them to colonize. However, our study verified that reducing the amount of phenolic acids in the rhizosphere, would attenuate the recruitment of soil-borne pathogenic fungi to to colonize in the rhizosphere.

### Notably, reducing rhizosphere metabolites would inhibit the synergistic damage of root secretions with pathogens

A growing body of research indicates that synergistic damage may be a significant driver of increased plant pests and diseases, chemosensory autotoxicity, and degradation of the soil environment. Phenolic acid significantly enhanced *Fusarium* damage to plants in a concentration-dependent manner, which increasing *Fusarium* acid secretion(22), disrupting host cell membranes and impairing mitochondrial function and metabolism, inhibiting defense enzymes and thereby reducing plant resistance, resulting in synergistic infestation with other pathogens, and exacerbating disease incidence(23). Meanwhile, *Fusarium oxysporum* evades the plant’s immune response and becomes an endophyte, promoting phenolic acid synthesis and release(24). This synergistic effect will attract more pathogens, cause root autotoxicity and soil acidification to suppress the growth of bacterial communities(25), and break the ecological balance of “plant-soil-microorganisms.”(26). Altogether, our study demonstrated that reducing rhizosphere metabolite declines phenolic acids with autotoxic effects, which may lower attraction to substrate-preferring pathogenic fungi, leading to diminished synergistic pathogenicity of root secretions and pathogens and thereby balancing the microbial ecology. Ultimately, rhizosphere metabolite modulation achieved a balanced microbial ecology with decreased fungal colonisation to the rhizosphere, particularly pathogens.

Reducing rhizosphere metabolites may prevent pathogens from accessing essential substrates. However, whether it could impact the microbial community, potentially harm the microbial ecology and overall health of the plant. Remarkably, our findings indicated the opposite outcome - a boost in bacterial diversity and improved plant resistance and growth.bacterial diversity and a more favorable plant resistance and growth. Our experiment showed that decreasing root secretion had a more significant impact on the fungal community’s structure, implying that rhizosphere metabolites are being utilized more by fungi or pathogenic fungi that occupy a significant ecological niche. Therefore, reducing rhizosphere metabolites may help regulate the fungal community structure and promote a balanced microbial ecology.

### Fungi can utilize macromolecules, while bacteria do not require excessive nutrients or root secretions distinct from fungi to regulate their communities

The alteration of fungal structure will help the bacterial community occupy the ecological niche, contributing to a shift from a “fungal” to a “bacterial” dominated soil ecosystem. The higher the bacterial diversity, the more favorable the ecological environment(27). The soil with higher microbial diversity tends to have more ecological functions and higher resistance to environmental stress. Communities with high diversity have high resource utilization complementarity to use resources in the environment entirely, which leaves fewer resources available for pathogens to invade(28), ultimately helping the ecosystem to remain healthy.

Reducing rhizosphere metabolites alleviate soil acidification and deterioration. The accumulation of root secretion directly leads to decreased soil permeability and pH and promotes the H^+^ efflux of pathogens to exacerbate soil acidification further(16), resulting in a suitable acidic environment for inhibiting beneficial and accumulating specific pathogens(29). However, regulating pH can help break this cycle, alleviate the continuous crop barrier, and increase bacterial diversity. capacity of soil microbiome to fight pathogenic Fusarium infections

In summary, our research highlights that rhizosphere metabolite plays a critical role in microbial ecology. Altering rhizosphere metabolites modulates microbial community composition, diversity, and function, vital in alleviating soil acidification, increasing bacterial diversity, and reducing the recruitment of fungi, especially pathogens, to the rhizosphere and exacerbating soil-borne disease. Therefore, it helps optimize the microbial ecology of successional soils and promote plant stress tolerance. At the same time, plant-microbiome interactions are neither beneficial nor detrimental but rather act as regulators, generating new phenotypes by reshuffling existing traits. The core microbiome is regulated and crucial in stress tolerance.

Within the core microbiota, such as ‘biomarker microorganisms,’ can influence the community structure through biotic solid ineractions with the host or with other microbial speciest(30), and that function as the first line of defense against pathogens, as their removal results in the loss of interactions-for example *Enterobacter cloacae*(31). We have also targeted the “biomarker microorganisms” and identification of their functions in microecology which played a more significant ecological role in optimizing the rhizosphere environment by efficiently utilizing soil nutrients, defending against pathogens, and participating in the metabolic function of reducing phenolic acid autotoxicity metabolites.

### However, explicit consideration of fundamental ecological processes for developing complex microbial communities is in its infancy

Microorganisms have long been applied as inoculants for biocontrol or biostimulation, their field efficacy varies with the soil environmental adaptability, but is critical to the rational design and manipulation of microbiomes in agricultural systems(9). In recent years, synthetic microbial communities (SynComs) of varied complexity have been constructed using bottom-up combinations and have been applied to plants as a means to study various aspects of plant microbiome interactions, including elucidation of the specific mechanisms that drive community assembly and the interactions among different members(32). Within this concept, functional keystone species can be predicted through topological networks derived from interactions and through metabolic models(33). This approach provides a pathway to maximize SynCom persistence and trait expression success in natural settings. It provides to understand how the ‘keystone behavior’ of identified hub species can be experimentally validated(34).

In conclusion, combining the results of our predictions of soil microbial community patterns used by omics-based approaches with the next step of actual microbial interactions used by SynComs-based approaches, we can gain a better elucidation of the relationship between microbial interactions and soil-borne disease and harness their beneficial properties against pathogens to optimize rhizosphere microbial ecology in a global ecosystem.

Moreover, why plants produce so many root metabolites that allow for autotoxins needs to be investigated, which goes against the natural plant survival laws. We hypothesize that it is the intensive agricultural model, which is anthropogenic. In the future, applying mathematical models of metabolic integration to synthetic microbial communities can better elucidate microbial interactions under metabolic regulation and the impact of microbial metabolites on the ecological functioning of the community.

## MATERIALS AND METHOD

**FIG7.**
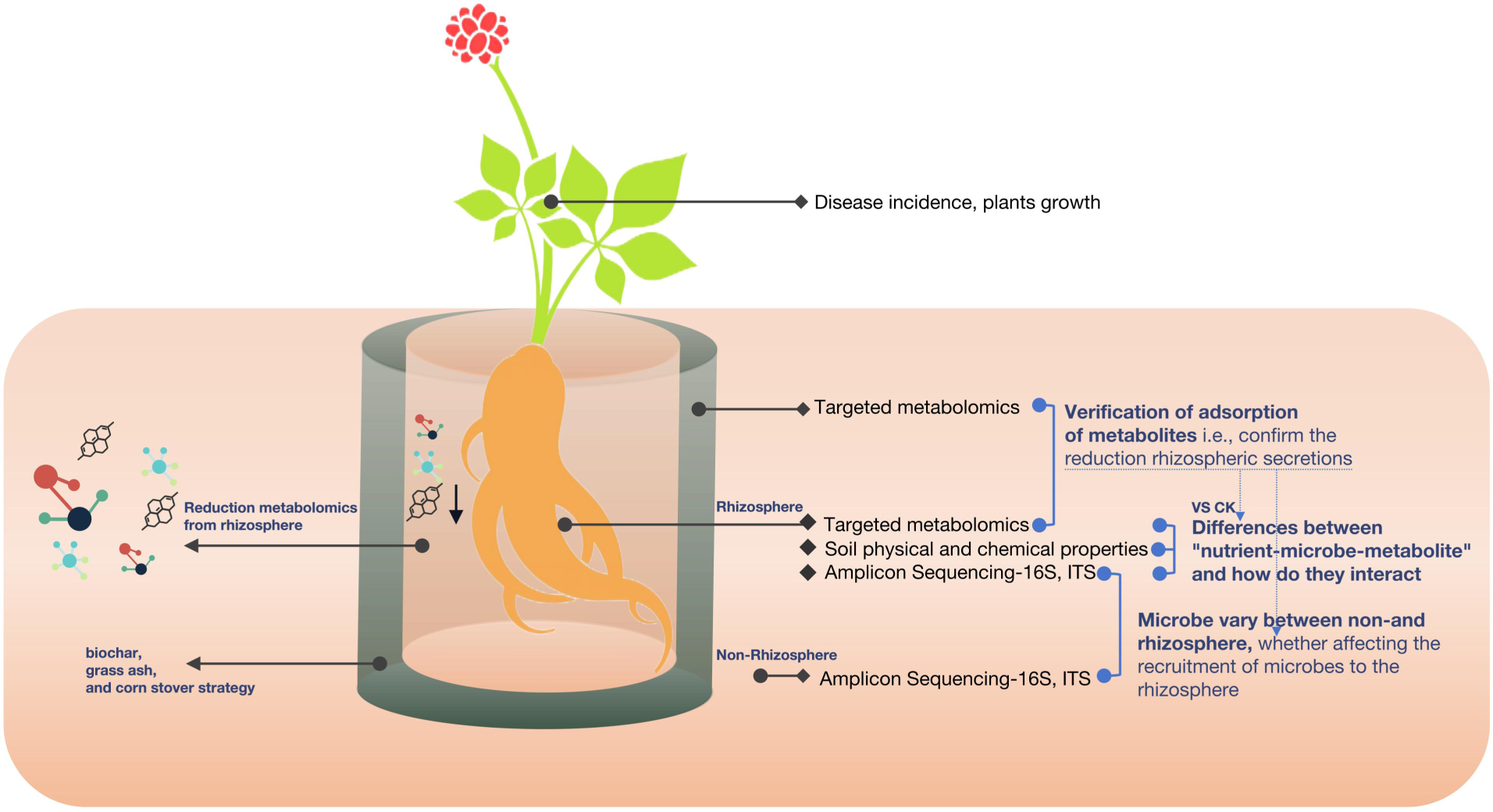
Reduce root secretion of materials and method of design map.

### Design of strategies to reduce root secretion. Growth, disease development, and soil collection

We selected *Panax ginseng* as the model plant, which has a lower survival rate than other crops due to continuous cropping obstacles caused by soil-borne fungal diseases. Therefore, We selected continuous cropping soil as the model soil. *Panax ginseng* seeds were selected to be robust, intact, and free from insect pests. We sterilized the seeds and seedling substrate, planted them in the substrate, and grew them into seedlings for a year by performing regular management. We decided to reduce metabolites from the rhizosphere by adsorption rather than adding soil conditioners directly to the soil rhizosphere, which would exclude the effects of other factors, and instead investigate the role of reduced metabolites on microbial ecology and plant resistance and growth. We chose three strategies shown to be able to adsorb metabolites, i.e., biochar, grass ash, and corn stover. We used the root bag method to experiment. Small and large bags were sewn with 25 μ m mesh diameter nylon mesh. The small bags were filled with *Panax ginseng* seedlings planted in the test succession soil. Three strategic substances were added into the large bag and the small bag with seedlings was placed into it, forming the interlayer containing three adsorption effects respectively. Then, The large bag was placed in a pot, and the outside was filled with the same soil (without the added substances). A control was set up where the interlayer was the same continuous soil. After a particular growth stage, we investigated the plants’ growth and seedling survival rates(33), removed the rhizosphere, non-rhizosphere, and interlayer, and set them aside.

### Profiling the non- and rhizosphere of Fungi and Bacteria

The sequencing was performed using the Illumina MiSeq, a third-generation high-throughput sequencing platform. Concentration of DNA was verified with NanoDrop and agarose gel. The genome DNA was used as template for PCR amplification with the barcoded primers and Tks Gflex DNA Polymerase (Takara).For bacterial diversity analysis, V3-V4 (or V4-V5) variable regions of 16S rRNA genes was amplified with universal primers 343 F and 798 R (or 515F and 907R for V4-V5 regions).For fungal diversity analysis, ITS I variable regions was amplified with universal primers ITS1F and ITS2.Raw sequencing data were in FASTQ format. Paired-end reads were then preprocessed using Trimmomatic software to detect and cut off ambiguous bases (N). It also cut off low quality sequences with average quality score below 20 using sliding window trimming approach. After trimming, paired-end reads were assembled using FLASH software. Parameters of assembly were: 10bp of minimal overlapping, 200bp of maximum overlapping and 20% of maximum mismatch rate. Sequences were performed further denoising as follows: reads with ambiguous, homologous sequences or below 200bp were abandoned. Reads with 75% of bases above Q20 were retained. Then, reads with chimera were detected and removed. These two steps were achieved using QIIME software (version 1.8.0). Clean reads were subjected to primer sequences removal and clustering to generate operational taxonomic units (OTUs) using Vsearch software with 97% similarity cutoff. The representative read of each OTU was selected using QIIME package. All representative reads were annotated and blasted against Silva database Version 138 (16s/18s rDNA) using RDP classifier (confidence threshold was 70%). All representative reads were annotated and blasted against Unite database (ITSs rDNA) using BLAST(35).

### Quantitative Wide-Target Metabolome Detection of the rhizosphere soil and interlayer metabolites

Sample preparation and extraction. After thawing the samples from the refrigerator at −80°C, mix them with vortex for 10 s.Take 2mL of the sample after mixing, place 2mL of the centrifuge tube, and then immerse the sample in liquid nitrogen as a whole. Put the sample into the lyophilizer for freeze-drying after the sample is completely frozen. After the samples were completely lyophilized, 200uL 70% methanol internal standard extract was added. Scroll for 15 min.iv Centrifuge (12000 r/min, 4 ° C) for 3 min. The supernatant was filtered with a microporous filter membrane (0.22 μm) and stored in a sample flask for LC-MS/MS test.

UPLC Conditions. The sample extracts were analyzed using an UPLC-ESI-MS /MS system (UPLC, SHIMADZU Nexera X2, https://www.shimadzu.com.cn/; MS, Applied Biosystems 4500 Q TRAP, https://www.thermofisher.cn/cn/zh/home/brands/ applied-biosystems.html). The analytical conditions were as follows, UPLC: column, Agilent SB-C18 (1.8 µm, 2.1 mm * 100 mm); The mobile phase was consisted of solvent A, pure water with 0.1% formic acid, and solvent B, acetonitrile with 0.1% formic acid. Sample measurements were performed with a gradient program that employed the starting conditions of 95% A, 5% B. Within 9 min, a linear gradient to 5% A, 95% B was programmed, and a composition of 5% A, 95% B was kept for 1 min. Subsequently, a composition of 95% A, 5.0% B was adjusted within 1.1 min and kept for 2.9 min. The flow velocity was set as 0.35 mL per minute; The column oven was set to 40℃; The injection volume was 4 μL. The effluent was alternatively connected to an ESI-triple quadrupole-linear ion trap (QTRAP)-MS(36).

ESI-Q TRAP-MS/MS. The ESI source operation parameters were as follows: source temperature 550°C; ion spray voltage (IS) 5500 V (positive ion mode)/-4500 V (negative ion mode); ion source gas I (GSI), gas II(GSII), curtain gas (CUR) were set at 50, 60, and 25 psi, respectively; the collision-activated dissociation(CAD) was high. Instrument tuning and mass calibration were performed with 10 and 100 μmol/L polypropylene glycol solutions in QQQ and LIT modes, respectively. QQQ scans were acquired as MRM experiments with collision gas (nitrogen) set to medium. DP(declustering potential) and CE(collision energy) for individual MRM transitions was done with further DP and CE optimization. A specific set of MRM transitions were monitored for each period according to the metabolites eluted within this period.

### Impacts of the rhizosphere microbial community function

For PICRUST analysis, we followed the suggested methods for OTU selection with Greengenes 13_ 5 using Galaxy (http://huttenhower.sph.harvard.edu/galaxy/). The predicted gene family abundances were analyzed using the Kyoto Encyclopedia of Genes and Genomes ethology group count level 3, and Storey FDR in STAMP software was used to avoid information of Type-I error. Carry out KEGG function prediction, enzyme classification number EC, and COG protein prediction, count the differences according to the Kruskal-Wallis algorithm, and select the different results to make a heatmap map.Use LEfSe(Linear differential analysis Effect Size) to display up/down-regulated function prediction and draw the differential heat map (37).

### Soil Physico-Chemistry Analysis

The content of organic matter is determined by the potassium dichromate volumetric method. After acid dissolution, total nitrogen was determined by SEAL-AA3 continuous flow analyzer, total phosphorus was determined by molybdenum-antimony colorimetry, and total potassium and available potassium were determined by flame photometer. The available nitrogen content adopts the alkali-hydrolyzed diffusion method, and the available phosphorus content adopts the sodium bicarbonate method. Determination of Cation exchange capacity(CEC) by neutral ammonium acetate method(38).

### Investigating potential interactions among biomarker, soil nutrients, metabolites and pathogens

Use LEfSe(Linear differential analysis Effect Size) to display up/down-regulated microorganisms and draw the differential genus heat map. To perform correlation and model prediction analysis, correlation analysis between biomarker microorganisms with other microorganisms, soil nutrients, phenolic acids, and pathogen. Using RDA analysis and mapping. Based on two (or three) histological quantitative files, draw a correlation chord diagram, a cluster heat map, and a network interaction diagram.

### Statistical Analysis

After data processing, the community composition histogram was drawn to show the community structure distribution; Evaluate the species richness and distribution uniformity in the sample, analyze the alpha diversity, and draw the boxplot plot. Based on the diversity index, calculate the different significance of the diversity index in different groups through the plot analysis (Wilcoxon algorithm). Microbial multivariate statistical analysis: calculate the difference species (OTU or phylum, genus and species level) between different groups through a statistical algorithm (Wilcoxon), display the up-regulated and down-regulated differential microorganisms through LEfSe (Linear differential analysis Effect Size), and draw the differential species heat map. For PICRUSt analysis, we followed the suggested methods for OTU selection with Greengenes 13_ 5 using Galaxy (http://huttenhower.sph.harvard.edu/galaxy/). After processing the data, a histogram was drawn to show the distribution of the community structure. The species richness and distribution uniformity in the sample were evaluated, and the alpha diversity was analyzed, which was then represented using a boxplot. Based on the diversity index, the significance of the diversity index was calculated in different groups through plot analysis (using the Wilcoxon algorithm).

## ACKNOWLEDGMENTS

This work was financially supported by the National Natural Science Foundation of China (31760360), Natural Science Foundation of Yunnan Province of China(202301 AT070007), and Natural Science Agricultural Joint Foundation of Yunnan Province of China(202101BD070001-078).

The authors declare no conflicts of interest.

---

**Supplementary fig1.**
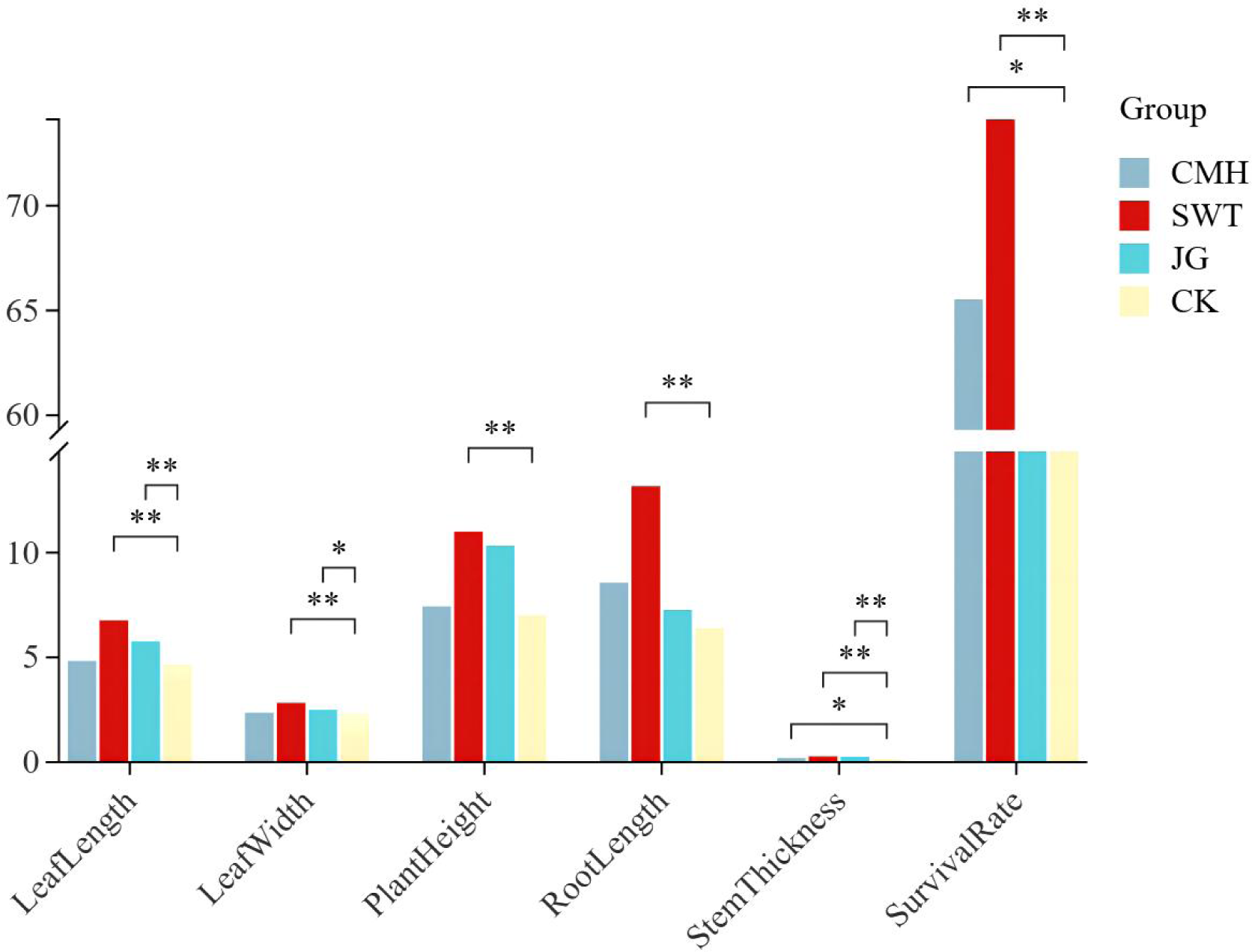
Plant growth and seedling survival in different strategies. CMH (grass ash), SWT (biochar), JG (corn stover), and CK (control).Plant growth and seedling survival were highest in the biochar strategy(*P*<0.01).

